# Single-protein diffusion in micrometer-size vesicles is governed by viscosity, surface charge, and membrane interactions

**DOI:** 10.64898/2026.10.21.760703

**Authors:** Jelmer Coenradij, Dmitrii Linnik, Kristine Postmus, Zeynap Şensan, Matilde Accorsi, Christiaan M. Punter, Rumiana Dimova, Bert Poolman

**Affiliations:** Department of Biochemistry, University of Groningen, Nijenborgh 3, 9747 AG, Groningen, Netherlands; Max Planck Institute of Colloids and Interfaces, Science Park Golm, 14476 Potsdam, Germany

**Author notes:** These authors contributed equally to this work (Jelmer Coenradij, Dmitrii Linnik).

**Keywords:** Giant unilamellar vesicles, protein mobility, single-molecule displacement measurement, membrane binding

## Abstract

Giant unilamellar vesicles (GUVs) are a powerful model system for studying protein dynamics in cell-sized compartments with a precisely tunable physicochemical environment. Here, we apply Single-Molecule Displacement Mapping (SMdM) to quantify how luminal viscosity, ionic strength, membrane lipid composition, and protein surface charge of soluble proteins govern intraluminal single-molecule diffusion. Increasing viscosity led to a predictable decrease in the diffusion coefficient. However, spatially resolved diffusion measurements reveal reduced protein mobility near the membrane, suggesting additional membrane-proximal interactions not captured by particle-based simulations. We observe long-range (∼1 µm) effects of membrane charge on protein mobility that extend well beyond classical Debye screening, which may reflect shape fluctuations of the GUV membrane. For the membrane-associated protein LactC2, we resolve two distinct diffusing populations within a single vesicle, corresponding to free and membrane-bound states. Our workflow provides a quantitative framework for probing single-protein mobility in vesicle-based synthetic cells. These results advance our understanding of how molecular crowding and transient membrane interactions shape protein diffusion within cell-sized compartments, with possible implications for metabolic organization.

## Introduction

Protein mobility is an important feature of living systems and provides insights into the spatiotemporal organization of the cell. A variety of microscopy techniques, including fluorescence recovery after photobleaching (FRAP), fluorescence correlation spectroscopy (FCS), single-particle tracking (SPT) and single-molecule displacement mapping (SMdM) are available to quantify the diffusion of molecules ^1,2^. Among these, SMdM offers the highest spatial resolution for measuring lateral diffusion coefficients *in vivo*, in both eukaryotic and prokaryotic cells^1,3^. Reconstructed diffusion maps can have a resolution < 50 nm, which allows detection of spatial heterogeneities in molecular mobility. Using SMdM, we previously demonstrated differences between apparent diffusion coefficients at the poles and mid-cell regions of *Escherichia coli*, as well as ageing-related changes in the dynamic structure of the cytoplasm^4,5^. SMdM has also been applied *in vitro* to study enhanced enzyme diffusion phenomena^6^ and surface aging of fused in sarcoma (FUS) protein condensates^7^.

Diffusion measurements can reveal details of the cellular organization, but data analysis is complicated by the presence of numerous (macro)molecules and active processes in the cell. Vesicle-based systems or synthetic cells offer a powerful alternative to overcome these challenges, because they are tunable in terms of concentration and composition of molecular species^8^. They can recapitulate life-like features, including compartmentalization, selective permeability and transport, information storage and processing, energy metabolism, and potentially growth and division^9^. Giant-unilamellar vesicles (GUV) are particularly attractive as cell-akin scaffolds because their micrometer size enables direct visualization of processes by optical microscopy and manipulation using microfluidic platforms^10–13^. In addition, GUVs allow vesicle-to-vesicle comparisons, facilitating systematic studies of size- and composition-dependent effects.

Synthetic cell research has focused on engineering modular functionalities^14^, such as lipid biosynthesis^15,16^, transcription and translation^15,17^, ATP production^18,19^, cell growth^20,21^, and celldivision^22,23^. In contrast, fundamental biophysical properties such as macromolecular crowding, intraluminal rheology and viscosity, and weak intermolecular interactions within synthetic cells remain comparatively underexplored^24,25^. Here, we present investigations aimed at elucidating the effect of viscosity, membrane charge, ionic strength and protein-membrane interactions on protein diffusion inside micrometer-sized vesicles.

We used SMdM to quantify the lateral diffusion coefficients of proteins. Specifically, we study the mobility of the fluorescent protein mEos3.2, two homologous TrxA- mEos3.2 fusion proteins that differ in surface charge^3^, and the LactC2-mEos3.2 fusion protein that binds to the membrane via the headgroup of the lipid phosphatidylserine (PS)^26,27^. These proteins are encapsulated in ∼20 µm GUVs composed of different mixtures of 1,2-dioleoyl-sn-glycero-3-phosphocholine (DOPC), 1,2-dioleoyl-sn-glycero-3-phospho-(1’-rac-glycerol) (DOPG), 1,2-dioleoyl-sn-glycero- 3-phosphoethanolamine (DOPE) plus 1,2-dioleoyl-sn-glycero-3-phospho-L-serine (DOPS), and varying luminal concentrations of potassium phosphate and/or sucrose. We observe a decrease in apparent diffusion coefficient *(D)* with increasing sucrose concentration (100 to 400 mM), in line with the Einstein-Stokes equation. We quantified the mobility of positively and negatively charged proteins as a function of distance from zwitterionic and negatively charged membranes and at varying ionic strength. We also measured the diffusion of LactC2-mEos3.2 in vesicles in which a fraction of the protein binds to PS in the membrane, and we observe a fast-diffusing population in the vesicle lumen and a slow-diffusing population at the membrane.

## Materials and Methods

### Chemicals

Luria Bertani broth (LB), Ampicillin, L-arabinose, DNase I, MgCl_2_, phenylmethylsulfonyl fluoride (PMSF), KCl, Glycerol, KH_2_PO_4_, K_2_HPO_4_, K-EDTA, Imidazole, polydimethylsiloxaan (PDMS), poly(vinyl alcohol) (PVA, fully hydrolyzed), 1,2-dioleoyl-sn-glycero-3-phosphocholine (DOPC) (Avanti), 1,2-dioleoyl-*sn*-glycero-3- phospho-(1’-rac-glycerol) (DOPG) (Avanti), 1,2-dioleoyl-sn-glycero-3- phosphoethanolamine (DOPE) (Avanti) and/or 1,2-dioleoyl-sn-glycero-3-phospho-L-serine (DOPS) (Avanti), BSA, Casein from bovine milk, Sucrose, Glucose

### Strains and plasmids

pBAD expression vectors, carrying genes encoding mEos3.2 and TrxA-mEos3.2 are from the Membrane Enzymology group^3^. The LactC2-mEos3.2 fusion construct was designed with mEos3.2 fused to the C-terminus of Lactadherin C2, followed by a 3C protease cleavage site and a C-terminal 10-histidine tag. The codon-optimized gene sequence was synthesized (GenScript, USA) and ligated in a pBAD expression plasmid via the Uracil-Specific Excision Reagent (USER) method (for primers see Supplementary Table S1). The resulting construct was transformed by heat-shock into *E. coli* MC1061 [F-, Δ(*araA-leu*)7697, Δ(*araD*139)*B/r*, Δ(*codB-lacI*)3, *galK*16, *galE*15(GalS), λ-, *e14-*, *mcrA*0, *relA*1, *rpsL*150(StrR), *spoT*1, *spoT*1, *hsdR*2] for gene expression.

### Gene Expression and Protein Purification

An overnight culture of *E. coli* MC1061 cells containing pBAD with the gene of interest (GOI) was grown in LB media supplemented with 100 µg/ml ampicillin. The following day, the culture was diluted 1:50 into fresh LB containing 100 µg/ml ampicillin and grown at 37 °C with shaking (180 rpm). Cell growth was monitored by measuring the OD_600_, and gene expression was induced when OD_600_ of 0.4-0.6 was reached by the addition of L-arabinose (0.01% v/v). The cultures were incubated for 4 hours at 37 °C with shaking (180 rpm). Next, the cultures were cooled on ice and cells were harvested by centrifugation (8000 x g, 20 min, 4 °C), followed by a washing step in 50 mM KPi pH 7.5, 200 mM KCl plus 10% v/v glycerol (buffer A) and collected again by centrifugation. The cell pellet was resuspended in the same buffer to a final OD_600_ of 150. DNase (100 µg/mL), MgSO_4_ (2 mM) and phenylmethylsulfonyl fluoride (PMSF, 1 mM) were added to the cell suspension prior to cell lysis via sonication (70% amplitude, 5 sec on, 10 sec off for a total time of 6 min). After sonication, 1 mM K-EDTA was added and cell debris was removed by centrifugation (42500 x g, 30 min at 4 °C).

For Immobilized Metal Affinity Chromatography (IMAC) purification, 5 mL of the cleared lysate (to which imidazole was added to a final concentration of 10 mM) was incubated with 1 mL Ni^2+^-Sepharose resin (Cytiva), preequilibrated with buffer A plus 10 mM imidazole, for 45 min at 4 °C with gentle nutation to allow protein binding to the resin. The flow-through was discarded, and the resin was washed with 20 column volumes (CV) of buffer A supplemented with 50 mM imidazole. Bound protein was eluted in 250 µl fractions using the same buffer but now containing 500 mM imidazole.

Protein concentrations were determined by measuring absorbance at 280 nm, using calculated extinction coefficients as given in Supplementary Table S2. Fractions containing the highest protein concentrations were pooled and concentrated to 10 mg/ml using centrifugal filter units (Amicon, 10 kDa MW cut-off). Next, K-EDTA was added to a final concentration of 5 mM, and the pooled sample was centrifuged (20,000 x g for 5 min, 4 °C) to remove potential aggregates.

Size-exclusion chromatography (SEC) was performed using a Superdex® 200 increase 10/300 column (Cytiva) equilibrated in 50 mM KPi pH 7.0, 100 mM KCl and 10% glycerol (buffer B). IMAC-purified protein was loaded onto the column and separated isocratically. Peak fractions were analyzed by absorbance at 280 and 505 nm, pooled, and concentrated to at least 10 mg/ml using centrifugal filter units (Amicon, 10 kDa MW cut-off). All purification steps were analyzed by SDS-PAGE.

For LactC2-mEos3.2, peak fractions obtained after SEC were pooled and incubated with 3C protease at a 1 to 5 (protease to LactC2-mEos3.2) ratio to cleave off the histidine tag. The mixture was nutated for 2 hours at 4 °C and subsequently subjected to another IMAC purification. Now, the resin was washed with 20 CV of buffer A supplemented with 30 mM imidazole. LactC2-mEos3.2 was eluted with buffer A supplemented with 80 mM imidazole. Elution peak fractions were pooled, concentrated and subjected to a final SEC purification step (same as described above).

### Microfluidic Chips Fabrication

The mask for the production of the microfluidic wafers was provided by Yandrapalli and Robinson^10^; the chip design, as described in Fig. 1 was reproduced on a silicon wafer through the exposure of SU-8 2035, a permanent negative epoxy photoresist purchased from micro resist technology GmbH (Berlin, Germany). Using this photoresist, the microfabrication process was done following the instructions provided in the SU-8’s technical data sheet.

**Figure 1.**
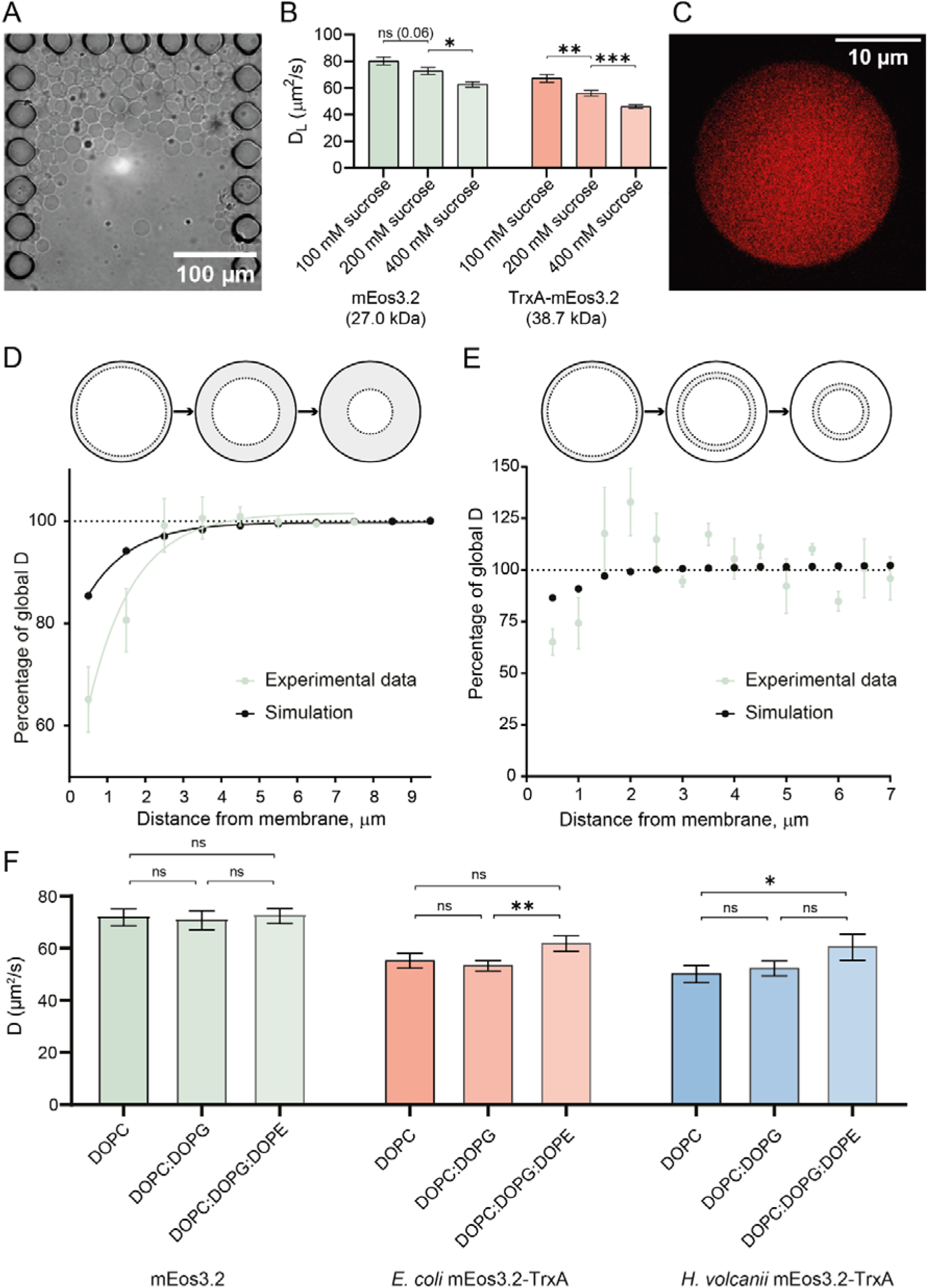
Diffusion coefficients of proteins encapsulated in GUVs composed of varying lipid compositions. (A) Representative bright-field microscopy image of the microfluidic chip bucket with trapped GUVs. GUVs were collected in a BSA- and Casein-passivated and buffer-equilibrated PDMS chip, using a flow rate of 0.5 µl/ml. Over time GUVs accumulate until enough are trapped, which is followed by washing of the GUVs with isosmotic medium (specified in the Methods section) prior to measuring diffusion by SMdM (B) The diffusion coefficients of mEos3.2 and *E. coli* TrxA-mEos3.2 in DOPC GUVs. The diffusion coefficients were determined by SMdM in GUVs containing 50 mM KPi pH 7.0 and varying concentrations of sucrose. Pairwise comparison was performed using t-test. Representative displacements fits are presented in Supplementary Figure S4. (C) Photoactivated localization microscopy (PALM) reconstructed image of 2 μM mEos3.2 encapsulated in GUVs, scale bar 10 µm. (D) The diffusion coefficient of mEos3.2 in DOPC GUVs decreases in close-to-border areas of the vesicle membrane (schematic of area selection for increment analysis on top of the graph). (E) The diffusion coefficient decreases in close-to-border rings of the vesicle membrane (schematic of area selection for ring analysis on top of the graph). The diffusion coefficient in 0.5 µm-thick “rings” at different distance from the GUV membrane. X-axis shows the distance from the middle of the “ring” to the border. Simulations show a confinement effect in rings closer than 1.5 µm to the border - similar distance as in the experimental data. (F). Average diffusion coefficients of mEos3.2 and mEos3.2-TrxA fusions in GUVs of different lipid compositions. mEos3.2 and the TrxA-mEos3.2 diffusion coefficients were determined by SMdM in GUVs containing 50 mM KPi pH 7.0, 200 mM sucrose plus 2 µM of protein. The exact diffusion coefficients, diameters and number of analyzed GUVs are presented in Supplementary Tables S3 and S5. Pairwise comparison was performed using t- test and exact p-values are presented in Supplementary Table S4. All data presented as mean ± SEM in the figure and as mean ± SD in Supplementary Table S4. Significance levels are presented as asterisk signs: (*) for p<0.05, (**) for p<0.01 and (***) for p<0.001.

The chips were fabricated according to the design and protocol established by Yandrapalli and Robinson^10^. Briefly, polydimethylsiloxane (PDMS) was prepared by thoroughly mixing 10:1 (w/w) SYLGARD^TM^ 184 Silicone Elastomer base and SYLGARD^TM^ 184 Silicone Elastomer Curing Agent. The mixture was degassed under vacuum for 30 min to remove air bubbles. A silicon wafer containing the microfluidic master mold was cleaned with a gentle stream of compressed air to remove dust particles and secured in a container. The degassed PDMS was carefully poured onto the wafer to form a layer of approximately 5 mm thick, followed by an additional 15 min of degassing. The PDMS was cured in an oven at 80 °C for 2 h. After curing, the PDMS layer was carefully peeled from the wafer and individual microfluidic chips were excised using a scalpel. Inlet and outlet holes were created using a 1.5 mm biopsy punch. 1.5H high-precision glass coverslips were cleaned by sonication in 5M KOH for 45 min, followed by bath sonication in milli-Q^TM^ water for 15 min. The coverslips were then thoroughly rinsed with milli-Q and dried under a stream of compressed air. Both the glass coverslips and PDMS chips were exposed to high oxygen plasma for 1 minute to activate the surfaces by making them hydrophilic allowing bonding. Immediately after plasma treatment, PDMS chips were irreversibly bonded to the cleaned glass coverslip. To create the inlet reservoir, a cut pipet tip was fixed over the inlet opening using residual uncured PDMS as an adhesive. The assembled devices were further cured at 80 °C for 30 min to ensure a secure seal.

The design of the produced chips is described in Figure 1 and the supplementary information of ref. 10: the design on the wafer provides 10 chips per chip production process. Each chip contains 12 channels, resulting in a more manageable distribution of GUVs in the chip; every channel contains 16 vesicle traps. Vesicle traps consist of 19 PDMS posts (diameter ∼37 μm) arranged in the shape of a 3-sided box, separated by ∼7 μm gaps to ensure solution flow.

### GUVs Formation by Gel-Assisted Swelling

GUVs were prepared using poly(vinyl alcohol) (PVA) gel-assisted swelling^28^. A 5% (w/v) PVA solution was prepared by dissolving PVA in milli-Q water while heating to 95 °C and continuous stirring until fully dissolved. Subsequently, 15 µl of the PVA solution was evenly spread onto two objective slides using a pipette tip. The coated slides were dried in an oven at 50 °C for 30 min to form a thin PVA gel layer.

Lipids were prepared separately. 1,2-dioleoyl-sn-glycero-3-phosphocholine (DOPC), 1,2-dioleoyl-*sn*-glycero-3-phospho-(1’-rac-glycerol) (DOPG), 1,2-dioleoyl-sn-glycero- 3-phosphoethanolamine (DOPE) and 1,2-dioleoyl-sn-glycero-3-phospho-L-serine (DOPS) were dissolved in chloroform at 25 mg/mL, mixed in the desired molar ratio and diluted to a final concentration of 4 mM in chloroform. Using a 25 µl Hamilton syringe, 5 µl of the lipid solution was spread evenly onto the dried PVA gel to form a homogeneous lipid film. The chloroform was removed by desiccation for 45 minutes.

A chamber was assembled by placing two lipid-coated slides face-to-face with a Teflon spacer in between to form a sealed ‘’sandwich’’ structure. A total volume of 700 µl swelling solution, containing the soluble components (50 mM KPi pH 7.0, 200 mM sucrose plus 2 µM of protein of interest, unless otherwise stated) was carefully injected in the chamber using a syringe equipped with a thin needle (0.4 mm diameter). GUVs were allowed to swell for 30-45 min at room temperature, ensuring the encapsulation of the swelling solution within the GUVs.

After swelling, the GUVs were harvested by gentle agitation of the chamber to detach the vesicles from the gel surface, and the swelling solution containing the vesicles was carefully collected from the chamber using a syringe equipped with a 0.4 mm needle. GUV formation was monitored using a table-top optical microscope (Leica).

### Single-molecule displacement mapping

#### Data Acquisition

Prior to imaging, GUVs were immobilized in a microfluidic chip. The microfluidic chips were passivated by adding 400 µl of 50 µM bovine serum albumin (BSA), followed by 400 µl of 2 mg/mL casein from bovine milk dissolved in mili-Q water via base addition followed by adjustment to pH 7.0 by acid titration, to the inlet reservoir. The chip was centrifuged using a 96 well plate platform centrifuge (900 x g, 6 min, 20 °C) to fill and coat the channels. After centrifugation, the chip was incubated with the casein solution for 10 min at room temperature. The microfluidic chip was mounted on the stage of Olympus IX-81 home-built wide-field microscope and the outlet was connected to a syringe pump (NE-1000 New Era SyringeOne). The remaining passivation solution in the reservoir was removed and the chip was flushed with at least 80 µL of washing buffer (50 mM KPi, pH 7.0 supplemented with glucose to match the osmolality of the vesicle lumen) at a flow rate of 10 µL/min. The flow rate was then reduced to 0.5 µL/min, and 50 µL of the GUV suspension was added to the inlet reservoir to allow vesicle trapping. After visual confirmation of trapped vesicles, the reservoir solution was replaced with washing buffer and flushed for 1h at 0.5 µL/min to remove background fluorescence originating from non-encapsulated protein. This flow rate was maintained for further measurements to prevent vesicles from moving.

SMdM measurements were performed as previously described^3,4^ with the following modifications: a 1 ms low power 405 nm laser pulse (OBIS 405 LX, 50 mW max. power) was used to photoconvert mEos3.2 from a green fluorescent state (507 nm ex. / 516 nm em.) to a red fluorescent state (572 nm ex. / 580 nm em.), and two 0.5 ms 85 mW readout beams of a 561 nm laser (OBIS LS 561-150) were used with time separation (Δt) of 1.5 ms. The emitted signal was collected by an EM-CCD camera (C9100-13, Hamamatsu), using an ET 605/70 M bypass filter (Chroma). The intensity of the 561 nm readout beams was reduced to 85 mW to minimize extensive photobleaching and high background fluorescence. For each measurement, 20,000 frames (∼10 min for acquisition time of 30 ms) were recorded for analysis.

#### Data Analysis

Single-protein localization, clustering and detection of displacements were performed as previously described^3,4^. The *r_max_* parameter, defining the maximal search radius for the displacements, was increased from 600 nm to 2 µm to account for the expected longer displacements. The diffusion coefficient of GFP in water^29^ was used to estimate the average protein displacement 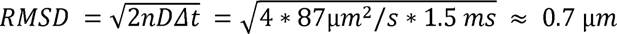, where n is dimensionality (n=2 in case of 2D diffusion).

Measured displacements were fitted using the adjusted probability density function (PDF) of a 2-dimensional random-walk diffusion model to obtain the diffusion coefficient:

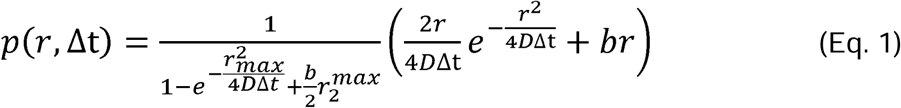

where *D* is a lateral diffusion coefficient, *r* is peak-to-peak displacement, Δ*t* is the time separation between 561 nm readout laser pulses, *b* is a background correction coefficient and *r_max_* is the maximum search radius (2 µm in this work). Equation 2 was used for the two-component diffusion model:

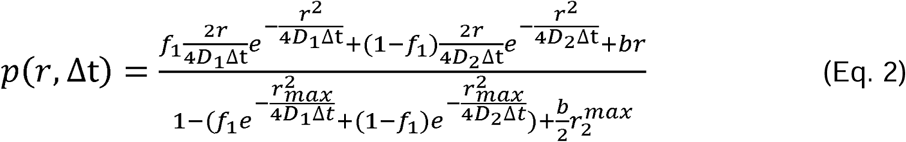

where *D*_1,2,_ are the diffusion coefficients and *f*_1_ is the fraction of the first diffusing population. The fraction of the second population is calculated based on *f*_2_ so that the total sum of fractions is 1. For analysis the fractions are converted to percentages.

To determine spatial variations in diffusion within individual GUVs (close to the border *versus* lumen), clustered fluorescence peaks from each circular GUV were enclosed within a circle of minimal radius representing the vesicle boundary, using Emo Welzl’s algorithm. Parameters such as circle center and radius were estimated. An “average” diffusion coefficient was first calculated using all displacements inside the vesicle.

To analyze close-to-border areas of different thicknesses we drew a concentric circle at the same central point but with radius smaller than the initial one by the given value (Figure 1D). We then selected different close-to-border areas with a step size of radius of 1 µm, staring from 0.5 µm away from the border, and calculated the ratio between the average diffusion coefficient and the diffusion coefficient calculated for the given close-to-border area. After plotting the calculated ratio as a function of the thickness of the analyzed area, we evaluate how the diffusion coefficient is decreased close to the lipid bilayer and how fast this decrease is compared to the average diffusion coefficient inside the GUV. This dependence can be fitted with equation *y = –*(*y_max_* – *y*_0_)*e^−kx^* for exponential plateauing, where *y_max_* is the plateau value, *y_0_*is the minimum *D* ratio, and *k* is an exponential increase factor. In the ring analysis (Figure 1E), we measured protein diffusion in separate “rings” of a fixed width of 0.5 µm, starting closest to the border of the GUV.

#### Particle diffusion simulation using Smoldyn

Particle diffusion simulations were performed using the Smoldyn software^30^. Smoldyn simulates random walk of particles by using an input diffusion coefficient and time steps to calculate each particle’s mean square displacement. A displacement is then randomly drawn from a normal distribution with variance corresponding to the mean square displacement. A random direction is independently chosen for each of the three spatial dimensions: x, y, and z. This procedure is repeated for every particle, as a result, each particle experiences a unique displacement and direction at every time step.

We simulated the diffusion in a spherical volume with elastic reflections from the border to evaluate the confinement effect on particle diffusion in GUVs. A total of 484 evenly distributed particles were placed inside a sphere with a diameter of 20 µm. Two simulations conditions were used with input diffusion coefficients of 55 µm^2^/s and 70 µm^2^/s to approximate the experimental diffusion coefficients for mEos3.2 and TrxA-mEos3.2, respectively. The total simulation time was 10 s with time steps of 1.5 ms, which matches the Δ*t* used in the SMdM experiments. For each condition five independent simulations were performed. Projected two-dimensional trajectories were obtained from the x and y coordinates of simulated particles within a 1 µm-thick equatorial section, corresponding to the approximate optical section thickness of the microscope. Displacements between consecutive frames were calculated and analyzed using the same pipeline as described for the SMdM experiments.

#### Photoactivated localization microscopy (PALM)

For super-resolution microscopy, the same home-built setup was used as for SMdM measurements. To convert and excite mEos3.2 fluorescent protein we utilized the same pulse pattern as for SMdM measurements. Briefly, a low-intensity 1 ms 405 nm photoactivation pulse was followed by a 0.5 ms 561 nm readout pulse within the same camera frame and again in the subsequent frame. This sequence was repeated for 20,000 frames. PALM reconstruction was then performed using the ThunderSTORM plugin [https://zitmen.github.io/thunderstorm/] of the ImageJ software as an Average Shifted Histograms visualization method.

### Statistical Analysis

Data in the text are presented as mean ± SD unless otherwise specified. Normality of data distribution was assessed using the Shapiro-Wilk test. Two-sided Student *t*-test was used at the significance level of 5 percent to compare mean values of two datasets. GraphPad Prism 8 software was used to perform all statistical tests and to make plots.

## Results

### Protein diffusion in GUVs is dependent of molecular weight and lumen viscosity

GUVs composed of DOPC were encapsulated with 2 µM mEos3.2 (M_W_ = 27.0 kDa, net surface charge is +2.21 e, Supplementary Table S2 and Supplementary Figure S1A) or TrxA-mEos3.2 fusion protein (M_W_ = 38.7 kDa, net surface charge is −2.48 e, Supplementary Table 2 and Supplementary Figure S1B), in the presence of 50 mM KPi pH 7.0 and increasing sucrose concentrations to modulate the lumen viscosity. Following GUV formation, the vesicles were immobilized in a PDMS-based microfluidic chip^10^ containing bucket-like trapping structures that facilitate vesicle accumulation, immobilization and controlled manipulation of the external medium. Non-encapsulated fluorescent protein was removed by washing with osmotically balanced buffer, to minimize background fluorescence. A representative brightfield image of trapped GUVs is shown in Figure 1A.

Apparent protein diffusion coefficients were determined by SMdM and the data are summarized in Figure 1B and Table S3A. As expected, the smaller mEos3.2 protein shows faster diffusion compared with the larger TrxA-mEos3.2 fusion protein. Increasing the internal viscosity resulted in a progressive reduction of the diffusion coefficients, a trend predicted by Einstein-Stokes equation. mEos3.2 in GUVs containing 100 mM sucrose displayed a diffusion coefficient of 80 ± 10 µm^2^/sec, comparable to GFP diffusion in aqueous buffer (87 ± 2 µm^2^/sec) as measured by FRAP^31^. To reduce protein mobility, internal sucrose concentrations were increased up to 400 mM, thereby increasing the internal viscosity. The minimum inter-frame interval of our microscope is 1.5 ms, which leads to relatively larger error in the apparent diffusion coefficient for faster diffusing proteins (Figure 1B). To slow-down diffusion while maintaining a good GUV yield, we used 200 mM sucrose in subsequent encapsulations. SMdM diffusion measurements with corresponding PALM reconstruction show that proteins distribute homogeneously in GUVs (Figure 1C), and the diffusion coefficient scales with the weight of the protein and the intravesicular viscosity (Figure 1B and Supplementary Table S3A).

### Protein Diffusion is Reduced in the vicinity of the GUV Membrane

SMdM has the unique feature to map how proteins move within the lumen of GUVs. Consistent with confinement effects^3^, protein diffusion is reduced near the membrane, which is visible on the reconstructed diffusion map (Supplementary Figure S2). To quantify the decrease in mEos3.2 diffusion, GUVs with an internal composition of 200 mM sucrose plus 50 mM KPi pH 7.0 were analyzed. First, an area 0.5 µm thick, adjacent to the membrane, was selected. Subsequently, the analyzed area was gradually extended to the center of the vesicle center in intervals of 1 µm (increment analysis). The relative diffusion coefficient (*D)*, expressed as percentage of the average diffusion coefficient (<*D>),* was determined as a function of distance from the membrane (Figure 1D). The reduction in mobility near the membrane was 65 ± 11% of <*D*>. At distances greater than 3.5 µm from the membrane, *D* converges to <*D>*.

As an alternative analysis, the vesicle interior was divided into concentric regions (ring analysis) of fixed width of 0.5 µm that were progressively displaced away from the membrane. The corresponding relative D was then compared to *<D>* (Figure 1E). Although this approach yielded larger fitting errors, because there is less data per ring, it also showed reduced mobility in the two outermost rings. Together, these analyses indicate that within approximately 1 µm from the membrane, the apparent protein mobility is decreased.

To benchmark the apparent confinement effects, particle diffusion was simulated using Smoldyn^30,32^. The simulations predict that proximity of a particle to a spherical compartment of similar size of the GUVs reduces the mobility to 85.4% ± 0.006% of <*D*> (Figure 1D,E and Supplementary Figure S3). In these simulations we used boundary with perfect elastic reflection of particles. Strikingly, this reduction is significantly less than the observed experimental reduction in diffusion coefficient. This indicates that geometric confinement alone cannot fully explain the reduced mobility near the membrane. Additional interactions, such as electrostatic interaction between the lipid bilayer and the proteins likely contribute to the pronounced decrease in diffusion at the membrane. Hydrodynamic coupling to the surface would also slow diffusion near a boundary and has been shown to affect protein mobility within ∼100 nm of supported bilayers^33,34^. For mEos3.2 with hydrodynamic radius ∼2.5 nm this effect would lead to at most ∼0.3% reduction in the diffusion coefficient at 1 µm from the surface, and therefore cannot account for the magnitude of the observed effect. Finally, the interaction of mEos3.2 with the lipids is most likely not elastic but a soft, dissipative reflection which may not be captured by Smoldyn simulations. Nevertheless, the distance from the GUV membrane at which protein mobility is decreased is in reasonable agreement with the Smoldyn confinement simulation.

### Effects of Membrane Surface Charge and Lipid Composition on Protein Diffusion

To investigate whether membrane surface charge contributes to the reduced protein mobility observed near the GUV membrane, we performed SMdM measurements of mEos3.2 (Supplementary Figure S1A), *E. coli* TrxA-mEos3.2 (Supplementary Figure S3B) and *H. vulcanii* TrxA-mEos3.2 (Supplementary Figure S1C); the M_W_ and net surface charge of the proteins is given in Supplementary Table S2. The proteins were encapsulated in neutral (zwitterionic) vesicles composed of 100 mol% DOPC and two types of negatively charged vesicles: DOPC:DOPG (75:25 mol%), and DOPC:DOPG:DOPE (50:25:25 mol%). The two negatively charged lipid compositions have same amount of charged DOPG lipid but differ in the amount and type of zwitterionic lipids (PE *versus* PC).

The average diffusion coefficient <D> of mEos3.2 did not differ significantly between the lipid compositions. Surprisingly, the <D> of *E. coli* and *H. vulcanii* TrxA-mEos3.2 were also not significantly different between DOPC and DOPC:DOPG vesicles, despite the large difference in net surface charge. The average diffusion coefficient of both TrxA-mEos3.2 fusion proteins was largest in DOPC:DOPG:DOPE vesicles (Figure 1F and Table S3B), for which we do not have an explanation.

Next, we analyzed the diffusion of the proteins as a function of distance from the membrane by the increment and ring analysis methods (Figure 1D and E; Figure 2). In DOPC:DOPG and DOPC:DOPG:DOPE vesicles, the mobility of positively-charged mEos3.2 decreased progressively toward the membrane, reaching approximately 12% ± 5% and 12% ±10% of *<D>* respectively. In contrast, in zwitterionic DOPC vesicles, mEos3.2 diffusion decreased substantially less, *i.e.* to 65 ± 11% of <*D*> (Figure 2A). For the moderately negatively charged *E. coli* TrxA-mEos3.2, diffusion in DOPC GUVs decreased to 35 ± 7% of *<D>,* whereas higher relative mobilities were observed in DOPC:DOPG (51% ± 19%) and DOPC:DOPG:DOPE (61 ± 5%) vesicles (Figure 2C). Similarly, the strongly negatively charged *H. vulcanii* TrxA-mEos3.2 exhibited a reduction to 24% ± 14% of *<D>* in DOPC vesicles, but only to 43 ± 10% and 41% ± 12% in DOPC:DOPG and DOPC:DOPG:DOPE vesicles, respectively (Figure 2 E).

**Figure 2.**
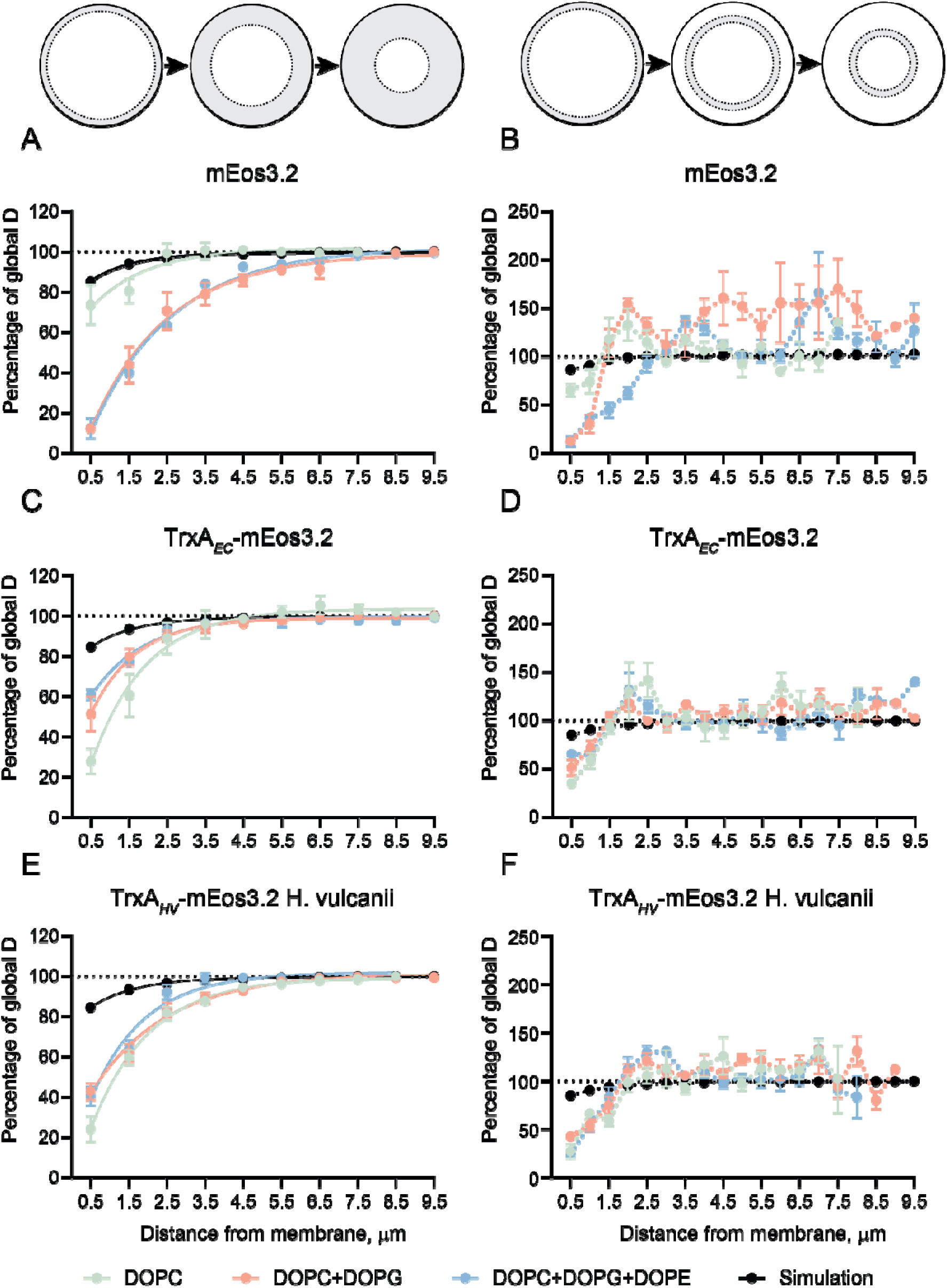
Reduction in diffusion coefficient close to the membrane depends on the charge of protein and lipid bilayer. Schematic of area selection for data analysis on top of the corresponding graphs. (A) Increment analysis of mEos3.2 diffusion. (B) Ring analysis of mEos3.2 diffusion. (C) Increment analysis of *E. coli* TrxA-mEos3.2 diffusion. (D) Ring analysis of *E coli* TrxA-mEos3.2 diffusion. (E) Increment analysis of *H. volcanii* TrxA- mEos3.2 diffusion. (F) Ring analysis of *H. volcanii* TrxA-mEos3.2 diffusion. Simulations predict a decrease up to ∼85% due to confinement effects for particles of sizes comparable with the analyzed proteins. For the increment analysis, the diffusion coefficient is measured in increasingly thick areas from the membrane; for ring analysis, the diffusion coefficient is determined in 0.5 µm-thick rings at different distance from the membrane. X-axis shows the thickness of the analyzed area or the distance from the middle of the ring to the membrane. Diameters and number of analyzed vesicles is presented in Supplementary Table S3; data presented as mean ± SEM and fit with exponential plateau equation.

Collectively, these data show a strong reduction in mobility of positively-charged mEos3.2 near negatively charged membranes, whereas the mobility of negatively-charged TrxA-mEos3.2 variants is comparatively enhanced under the same conditions. Ring analysis does not show changes in the distance from the membrane at which the protein mobility is affected by the confinement, so the observed charge-dependent effects are happening at relatively large (∼1 to 1.5 µm) distance from the membrane (Figure 2B,D,F).

### Modulation of Protein Diffusion by Ionic Strength

We then investigated the effect of ionic strength on the diffusion of the proteins. GUVs composed of DOPC were prepared in 10 mM, 50 mM (standard condition), and 200 mM KPi, pH 7.0 supplemented with 200 mM sucrose, corresponding to ionic strength values of 17.6, 88.1 and 353 mM, respectively. We hypothesized that lower ionic strength would enhance membrane charge effects because the ionic strength is reduced, and *vice versa* at high ionic strength. The average diffusion coefficients <*D*> of *H. volcanii* TrxA-mEos3.2 decreased progressively with increasing KPi concentration, which may reflect the increase in bulk solution viscosity^35,36^. A similar trend was observed for *E. coli* TrxA-mEos3.2 when the salt concentration was increased from 10 and 50 mM KPi, but <*D*> did not decrease further at 200 mM KPi (Figure 3A and Supplementary Table S3C).

**Figure 3.**
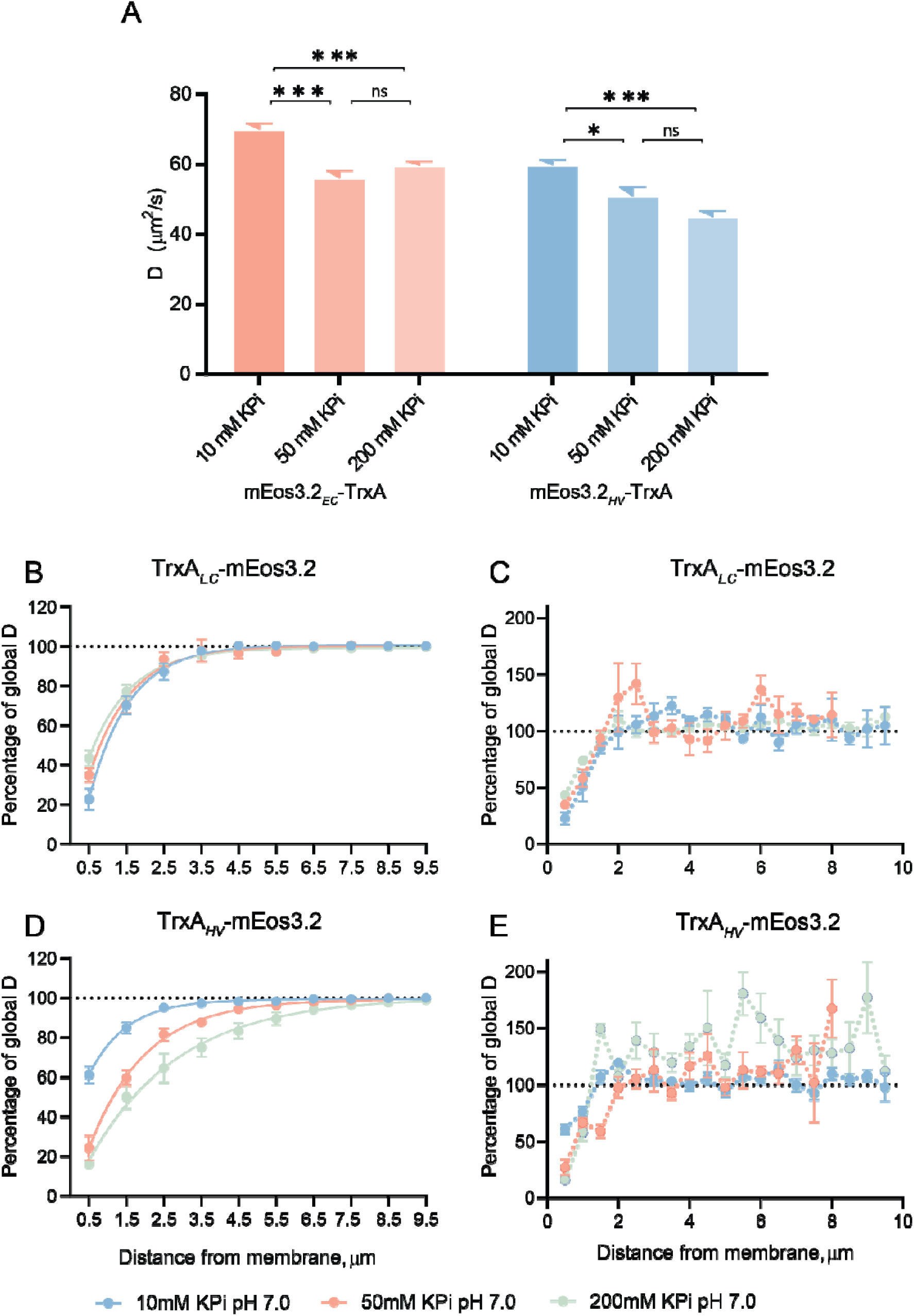
Diffusion of *E. coli* TrxA-mEos3.2 and *H. volcanii* TrxA-mEos3.2 at different salt concentrations. (A) Average diffusion coefficients of the TrxA fusion proteins in different buffer conditions. The exact diffusion coefficients, diameters and number of analyzed GUVs are presented in Supplementary Table S3C. Pairwise comparison was performed using t-test and exact p-values are presented in Supplementary Table S4. All data presented as mean ± SEM in the figure and as mean ± SD in Supplementary Table S4. Significance levels are presented as asterisk signs: (*) for p<0.05, (**) for p<0.01 and (***) for p<0.001. (B) Spatial diffusion by increment (left) and ring (right) analysis of *E. coli* TrxA- mEos3.2. (D) Spatial diffusion by increment (left) and ring (right) analysis of *H. volcanii* TrxA- mEos3.2. Diameters and number of analyzed GUVs are presented in Supplementary Table S5.

For near-membrane diffusion we observed protein-specific effects. For *E. coli* TrxA- mEos3.2 (net charge of −2.48), the effects of ionic strength were small but the trends were consistent with electrostatic screening (Figure 3B). In contrast, *H. volcanii* TrxA- mEos3.2 (net charge of −13.47) displayed a markedly different dependence on ionic strength. The diffusion near the membrane was reduced most (16% ± 5% of <*D*>) at 200 mM KPi, followed by 24% ± 14% at 50 mM KPi. Surprisingly, the mobility was least affected at 10 mM KPi, remaining at 61% ± 11% of <*D*> (Figure 3D). In the ring analysis, the diffusion coefficients converge to *<D>* at ∼1.5 µm from the membrane for both *E. coli* and *H. volcanii* TrxA-mEos3.2 (Figure 3C and E). The increment analysis yields similar recovery distances for *E.coli* TrxA-mEos3.2, but for *H. volcanii* TrxA-mEos3.2 the distances are much larger: 3.5, 6 and 9.5 µm for 10, 50 and 200 mM KPi respectively.

### Diffusion of Membrane-Associated Proteins

After characterizing non-specific, charge-dependent effects on the diffusion of soluble proteins, we investigated the mobility of mEos3.2-LactC2. mEos3.2-LactC2 (M_W_ = 45.35 kDa, net surface charge is +5.02, Supplementary Table S2 and Supplementary Figure S1D) contains the phosphatidylserine (PS) - binding domain C2 of lactadherin^26,27^ fused to the photoactivatable mEos3.2. GUVs were initially prepared with a lipid composition of DOPC:DOPS (75:25 mol%) in 50 mM KPi pH 7.0, 200 mM sucrose plus 2 µM encapsulated mEos3.2-LactC2 (about one mEos3.2-LactC2 per 100 PS). However, under these conditions, diffusion measurements were not possible because essentially all LactC2 was bound to the membrane, preventing detection of individual fluorescent peaks by SMdM.

To mitigate these effects, the lipid composition was adjusted to DOPC:DOPS (99.5:0.5 mol%, corresponding to about one mEos3.2-LactC2 per two PS in one leaflet for a vesicle of 20 µm in diameter), which improved the quality and integrity of the vesicles, however, LactC2 remained predominantly membrane-bound. Individual proteins could be detected, but diffusion measurements were not possible due to high local density of mEos3.2-LactC2 (Figure 4A). We then reduced the DOPS content to 0.01 mol% and lowered the mEos3.2-LactC2 concentration to 0.2 µM. The majority of LactC remained membrane-associated, even though the ratio of mEos3.2- LactC2 over inner leaflet DOPS was now 2.5:1 (protein:lipid). This microscopy observations make sense because membrane-bound LactC2 is concentrated locally, and free LactC is diluted over the entire GUV volume.

**Figure 4.**
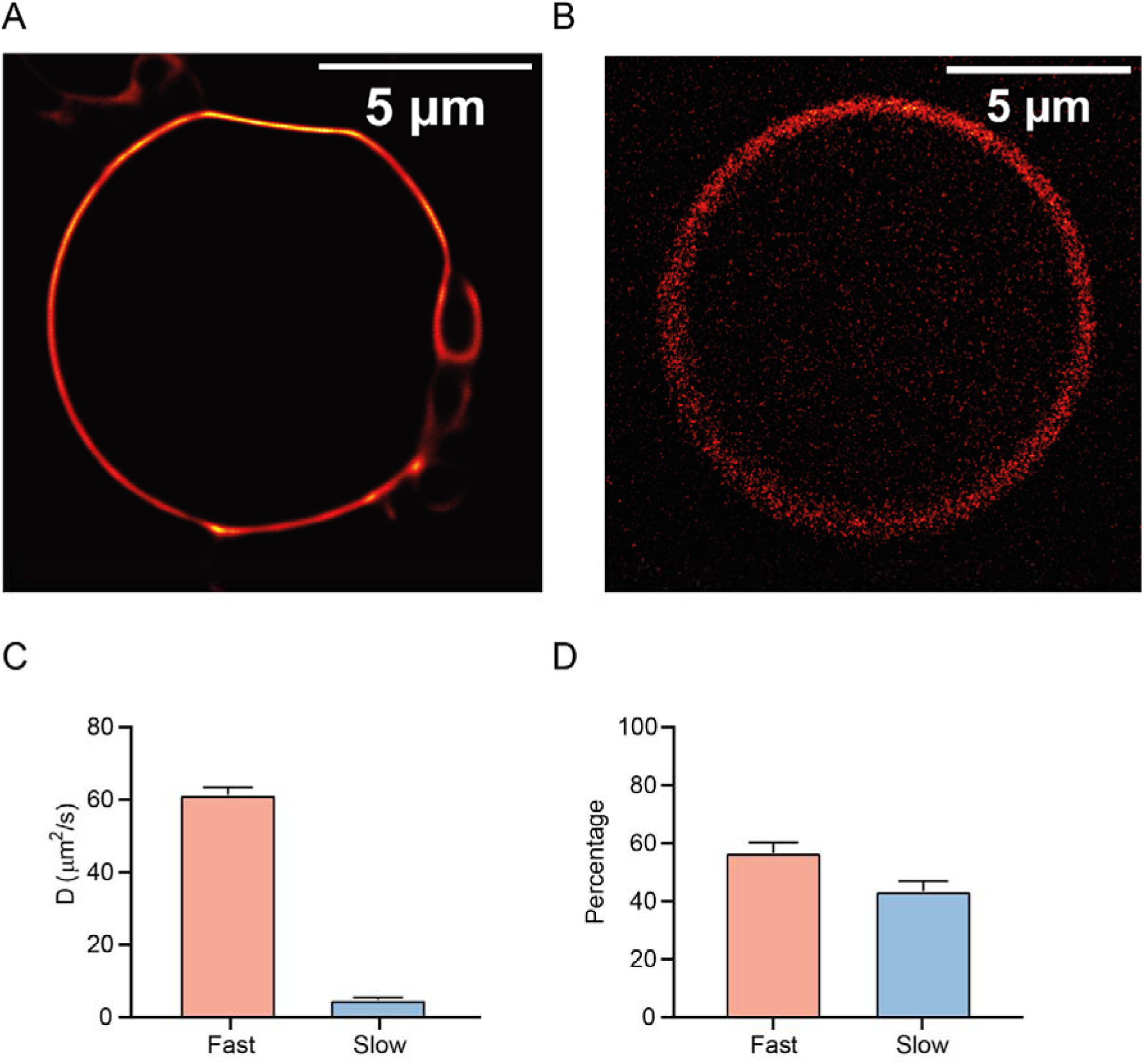
Effect of DOPS concentration on membrane binding and diffusion of LactC2- mEos3.2. Photoactivated localization microscopy (PALM) reconstructed image of the vesicles with different DOPS content. (A) DOPS content at 0.5 mol% results in complete association (sticking) of LactC2-mEos3.2 (2 µM) to the membrane. (B) DOPS content at 0.001 mol% resulted in conditions suitable for SMdM measurements of LactC2-mEos3.2 (0.2 µM). (C) LacTC2-mEos3.2 diffusion coefficients measured in vesicles from panel B. In this condition, we observed a fast- and slow-diffusing component. Representative displacements fits are presented in Supplementary Figure S5 (D) The percentages of the single-protein displacements associated with fast and slow diffusing molecules; the exact values, SEM and number of replicates are indicated in Supplementary Table S3D. For the SMdM measurements only non-deformed GUVs (0.001 mol% of DOPS) were used.

Only upon reducing the DOPS fraction to 0.001 mol% (about one mEos3.2-LactC2 per 0.04 PS) we could observe by SMdM membrane-bound and freely diffusing mEos3.2-LactC2 (Figure 4B using equation 2). The fast-diffusing population had a <*D*> of 59.9 ± 6.9 µm^2^/sec, and the slow-diffusing population a <*D*> of 4.3 ± 2.0 µm^2^/sec, which is close to the diffusion coefficient of lipids in membranes^37^ (Figure 4C and Supplementary Table S3D). The fast component (freely diffusing mEos3.2- LactC2) amounted to 57% of the measured displacements, and the slow component (membrane-bound protein) to 43 % (Figure 4D).

## Discussion

Giant-unilamellar vesicles (GUVs) are great platforms to study complex cellular processes *in vitro* in a tunable cell-mimicking environment, applicable to various types of reaction and signaling networks, protein synthesis, and cell division^15,23^. Particle mobility influences the subcellular localization and interactions between proteins and other macromolecules^38–43^. In *vivo* this mobility is influenced by (macro)molecular crowding, rheology and viscosity of the intracellular environment, and interactions with cellular structures such as the membrane or DNA^44^. These interactions form the basis for the spatiotemporal organization of *e.g.* the cytoplasm of a cell. Here, we present a workflow to measure and quantify single-protein diffusion in GUVs using single-molecule displacement mapping (SMdM). For data acquisition, GUVs were immobilized in a microfluidic chip, enabling removal of non-encapsulated (fluorescent) components, controlled buffer exchange in isosmotic conditions, and allowing external medium perturbations. Below we discuss the limitations and advantages of our microscopy and analyses methods.

Apparent diffusion coefficients of mEos3.2 (27.0 kDa), and *E. coli* TrxA-mEos3.2 (38.7 kDa) were measured in DOPC GUVs while varying the internal sucrose concentration to change luminal viscosity or salt concentration to change the internal ionic strength. In our microscopy setup, the minimal time separation between the readout laser pulses is 1.5 ms. For diffusion coefficients in the range of 50 to 80 μm^2^/s, average molecular displacements are in the range of 0.6 to 0.7 μm, consistent with root-mean-square displacements predictions. Consequently, it is hard to robustly fit the displacement data with a probability density function of a 2D random walk model, i.e. if the displacements overlap with the noise, which is more pronounced at longer displacements (Equation 1). Increasing lumen viscosity, reduces the displacement length and thus improved fitting accuracy. However, at 400 mM sucrose, it became challenging to prepare GUVs by the gel-assisted swelling method. Hence, 200 mM sucrose was used in the majority of the experiments as a compromise between measurement accuracy and vesicle stability.

Spatially resolved diffusion analysis shows reduced mobility near the lipid bilayer. The ability to select areas within a GUV from a single experiment is a key advantage of SMdM over other microscopy techniques. The apparent diffusion coefficient <*D>* was calculated based on the displacements from the entire vesicle, and within progressively thicker close-to-membrane regions (increment analysis). The decrease of apparent mEos3.2 diffusion coefficient close to the DOPC membrane exceeds that predicted by particle-based simulations, while the distance at which its mobility starts to decrease is similar (Figure 2A). Alternatively, we analyzed *D* in concentric rings with 1 µm thickness, positioned at varying distance from the membrane. This independent analysis yielded a similar behavior, confirming that reduced mobility starts around 1.5 µm from the membrane.

In the increment analysis, short displacements are used for fitting data further from the membrane (Figure 3A,C,E), meaning that this analysis reflects relative rather than absolute increases in the diffusion coefficient with increasing distance from the membrane. In the concentric ring analysis (Figure 2B,D,F), we use only the displacements from a given small area and these plots are more representative of the actual diffusion, and the diffusion coefficient of rings inside GUVs is not biased by short displacements closer to the membrane (Figure 2B,D,F). Here, the values from the rings relatively far from the membrane have a value higher than <*D>*; the error is larger because there are fewer displacements than in close-to-membrane regions.

Diffusion of differently charged proteins was strongly affected by the surface charge of the membrane. Based on the net surface charges of the proteins (Table S2), we expected different behavior near membranes for the anionic and cationic proteins. Both increment and ring analyses showed a pronounced decrease of *D* for positively-charged mEos3.2 near negatively charged DOPC:DOPG and DOPC:DOPG:DOPE membranes. Negatively-charged TrxA-mEos3.2 in GUVs with 25 mol% anionic lipid (DOPC:DOPG or DOPC:DOPG:DOPE) showed no difference in diffusion behavior when compared to zwitterionic DOPC vesicles. Our data can be partially explained by electrostatic interactions between charged proteins and membranes. However, the distances over which we observe the effects extend over micrometers – far beyond nanometer-scale electrostatic distances (< 10 Å^45^). This suggests that factors other than electrostatics are affecting the diffusion near the membrane. One possible explanation involves the mechanical properties of the membrane such as membrane tension and bending rigidity^46^. These properties influence the amplitude and spatial extent of thermal fluctuations which could alter the local environment near the membrane and thus affect protein mobility through transient protein–membrane interactions^47^. Additionally, differences in lipid composition and surface charge may influence membrane mechanics and thus contribute to the observed differences in diffusion behavior^48^. Although the precise contribution of these effects remains unclear, our findings suggest that protein diffusion near lipid membranes may be influenced by a combination of electrostatic interactions and membrane mechanical properties.

Using SMdM, we were able to resolve the diffusion properties of membrane-bound and soluble LactC2-mEos3.2 (Equation 2). We had to keep the DOPS content of the membrane at or below 0.001 mol% DOPS (corresponding to an approximate DOPS:LactC2 ratio of 1:25) in order to avoid oversaturating the membrane with fluorescent protein. The diffusion coefficient of the fast LactC2-mEos3.2 fraction was 59.9 ± 6.9 µm^2^/sec, which is in line with the mobility of water-soluble protein with a molecular mass of about 45 kDa. The diffusion coefficient of the slow fraction was 4.3 ± 2.0 µm^2^/sec, which is in line with diffusion coefficients found for membrane-surface bound proteins^49^; the value is also similar to that of the lateral diffusion coefficient of DOPS in GUVs^37^.

The order of magnitude reduction in mobility is consistent with diffusion in a two-dimensional system (attached to the lipid bilayer) compared to three-dimensional (luminal volume of the GUV). The ability to distinguish these two diffusion regimes within the same vesicles highlights the power of SMdM for resolving distinct mobility states without requiring prior spatial selection. Together, these findings show that diffusion analysis provides a quantitative and sensitive readout of membrane-associated proteins, and they establish SMdM as a powerful tool to study peripheral membrane-protein interactions in minimal synthetic cell systems.

In summary, we established a pipeline to quantify single-protein diffusion in GUVs using SMdM. This approach enables systematic evaluation of how viscosity, lipid composition, membrane charge and composition, and ionic strength influence the mobility of soluble and membrane-associated proteins differing in size and net charge.

Our results demonstrate that diffusion in cell-sized compartments is governed not only by bulk viscosity but also by membrane-mediated effects. The ability to resolve multiple diffusion regimes within the same vesicle highlights the strength of SMdM for dissecting protein mobility in synthetic cell systems. The insights obtained by quantitative measurements of protein diffusion are essential for understanding how the physical environment of a cell shapes protein function, and for designing synthetic cells with programmable behavior.

## Description of supporting material

Supplementary Figure S1. SDS-PAA gel analysis of the purified proteins.

Supplementary Figure S2. Reconstructed diffusion map of mEos3.2 encapsulated in DOPC GUV

Supplementary Figure S3 Confinement simulations using Smoldyn software

Supplementary Figure S4. Representative probability density function fitting profiles of the mEos3.2 and TrxA-mEos3.2 displacements in DOPC GUVs.

Supplementary Figure S5. Two representative probability density function fitting profiles of the LactC2-mEos3.2 in DOPC:DOPS (99.99:0.001 mol%) GUVs.

Supplementary Table S1. Primers for USER cloning of LactC2-mEos3.2.

Supplementary Table S2. Parameters of the proteins used in this work.

Supplementary Table S3. Exact diffusion coefficients measured in this work.

Supplementary Table S4. Summary of p-values in this work.

Supplementary Table S5. Size of GUVs, used for close-to-border and “ring” areas analysis.

## Acknowledgements

We would like to thank Jeroen Niekamp and Amber van Doorn for helping with setting up the GUV immobilization and protein purification protocols. We are thankful to Ivan Maslov, Chrisopher Aberg, and Lyan van der Sleen for discussion of the data and/or commenting on the manuscript. The work of Dmitrii Linnik and Bert Poolman was funded by the NWO National Science Program “The limits to growth” (grant number NWA.1292.19.170) and ERC Synergy program grant MetaDivide. The work of Jelmer Coenradij and Bert Poolman was supported by the NWO Gravitation program “Building a Synthetic Cell” (BaSyC). Rumiana Dimova and Bert Poolman acknowledge the ComeInCell network (funded by the European Union’s Horizon Europe research and innovation program under the Marie Skłodowska-Curie grant agreement No. 101168939) which provided favorable interaction environment.

## Author contributions

J.C. and D.L. conceptualized, designed, executed and supervised the experiments. J.C. expressed and purified the proteins, prepared the GUVs and made the microfluidic chips. D.L. performed the microscopy experiments and data analysis. K.P., Z.S. executed the experiments. C.M.P and D.L. developed the software for data analysis. M.A. produced the wafers for microfluidic chips under supervision of R.D. B.P. conceptualized and supervised the research, and acquired funding. J.C., D.L. and B.P. wrote the manuscript with the help of all authors.

