## Supplementary materials for "Single-protein diffusion in micrometer-size vesicles is governed by viscosity, surface charge, and membrane interactions"

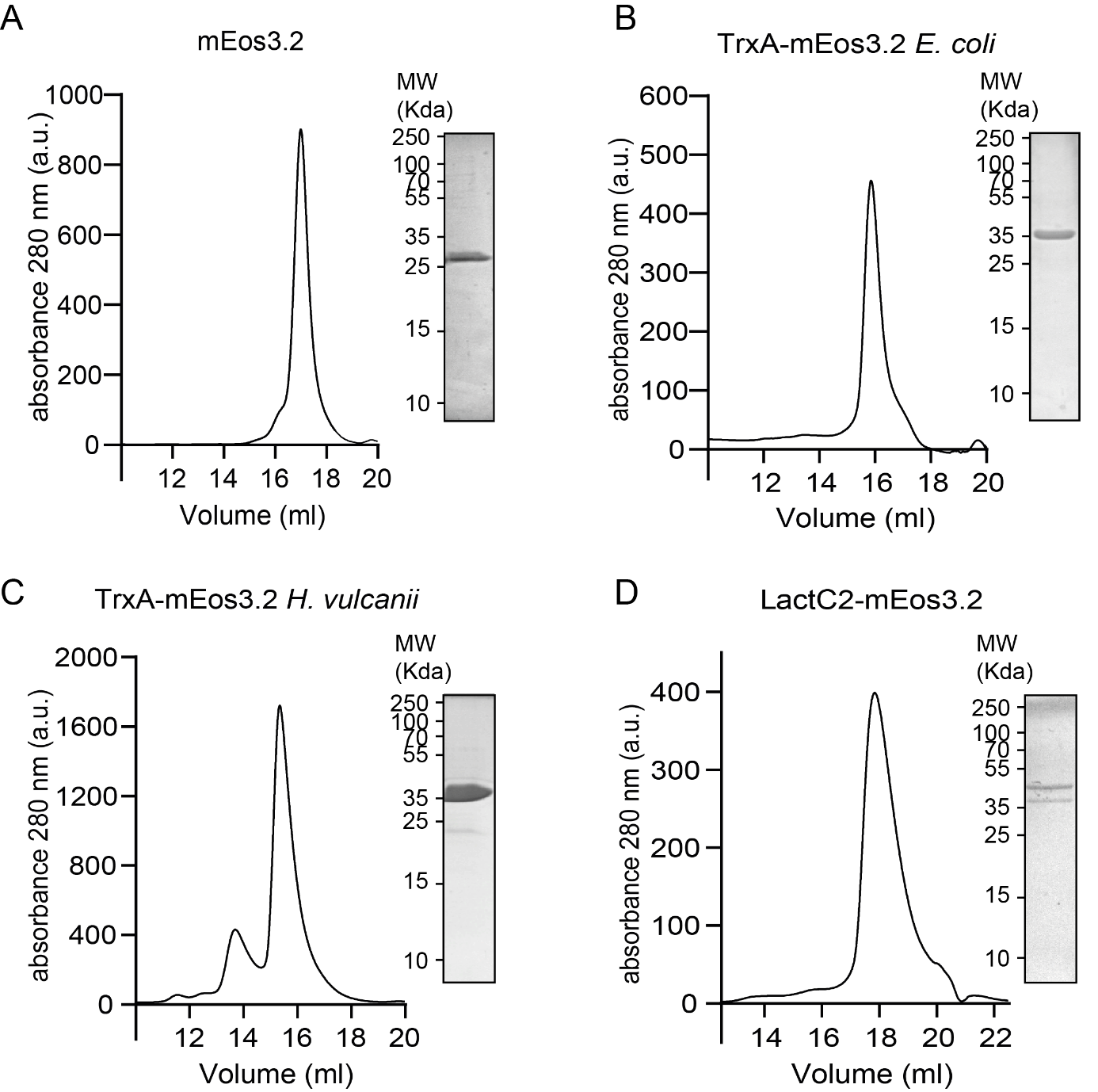

**Supplementary Figure S1. SDS-PAA gel analysis of the purified proteins**. All proteins were purified using IMAC followed by SEC (Superdex® 200 increase 10/300 column (Cytiva) chromatography. (**A)** mEos3.2 displayed a small shoulder preceding the main SEC elution peak. (**B)** TrxA-mEos3.2 from *E. coli* exhibited a small shoulder following the main elution peak. (**C)** TrxA-mEos3.2 from *H. volcanii* showed an additional peak preceding the main elution peak, which could be the dimeric state. (**D)** LactC2-mEos3.2 showed a minor shoulder on the right side of the main elution peak. SDS-PAA analysis (on the inset of each SEC profile) shows the purity of proteins.

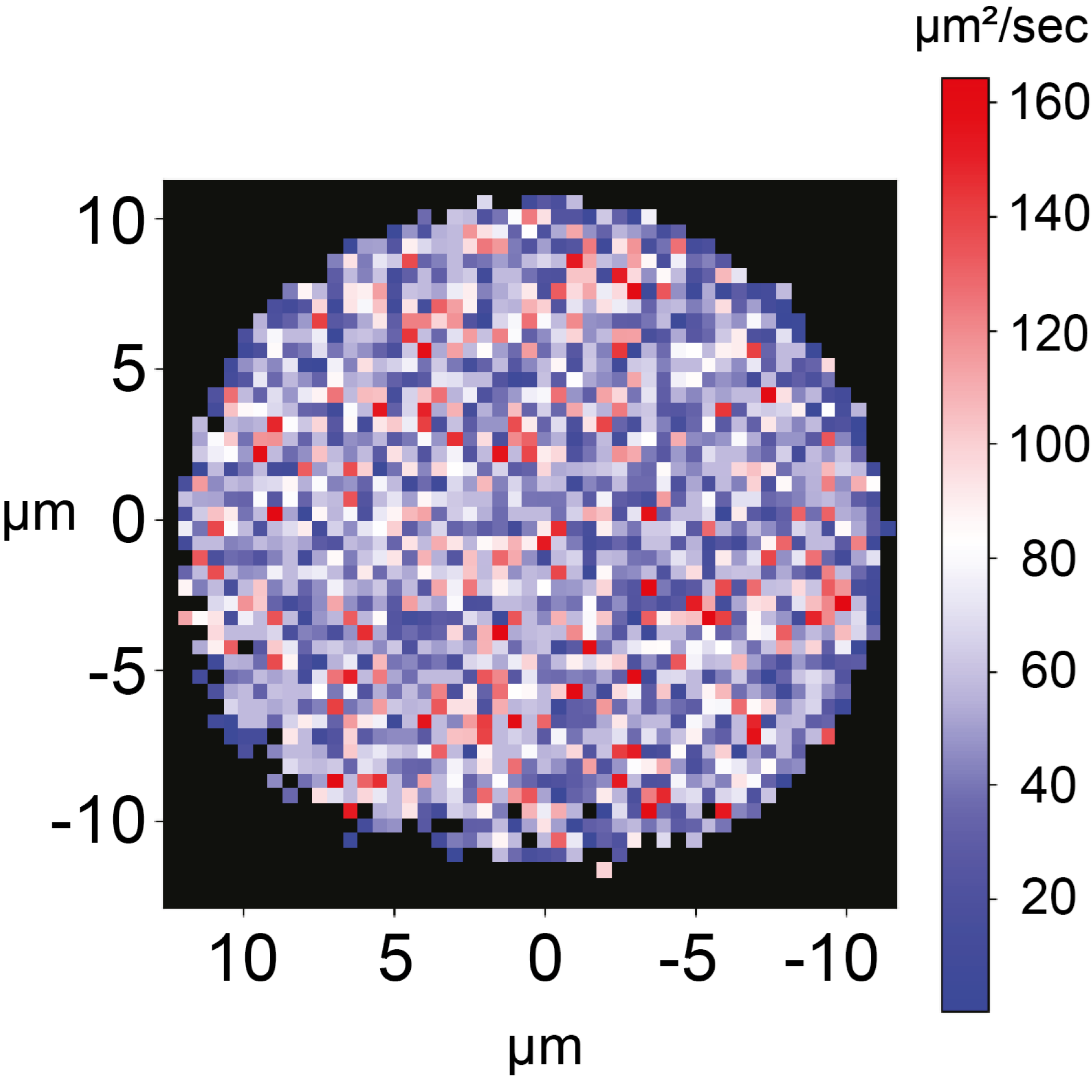

**Supplementary Figure S2. Reconstructed diffusion map of mEos3.2 encapsulated in DOPC GUV.** The internal medium was composed of 50 mM KPi pH 7.0, 200 mM of sucrose plus 2 µM of mEos3.2. Bin size = 0.5 µm, and displacements starting inside each bin were fitted with Equation (1).

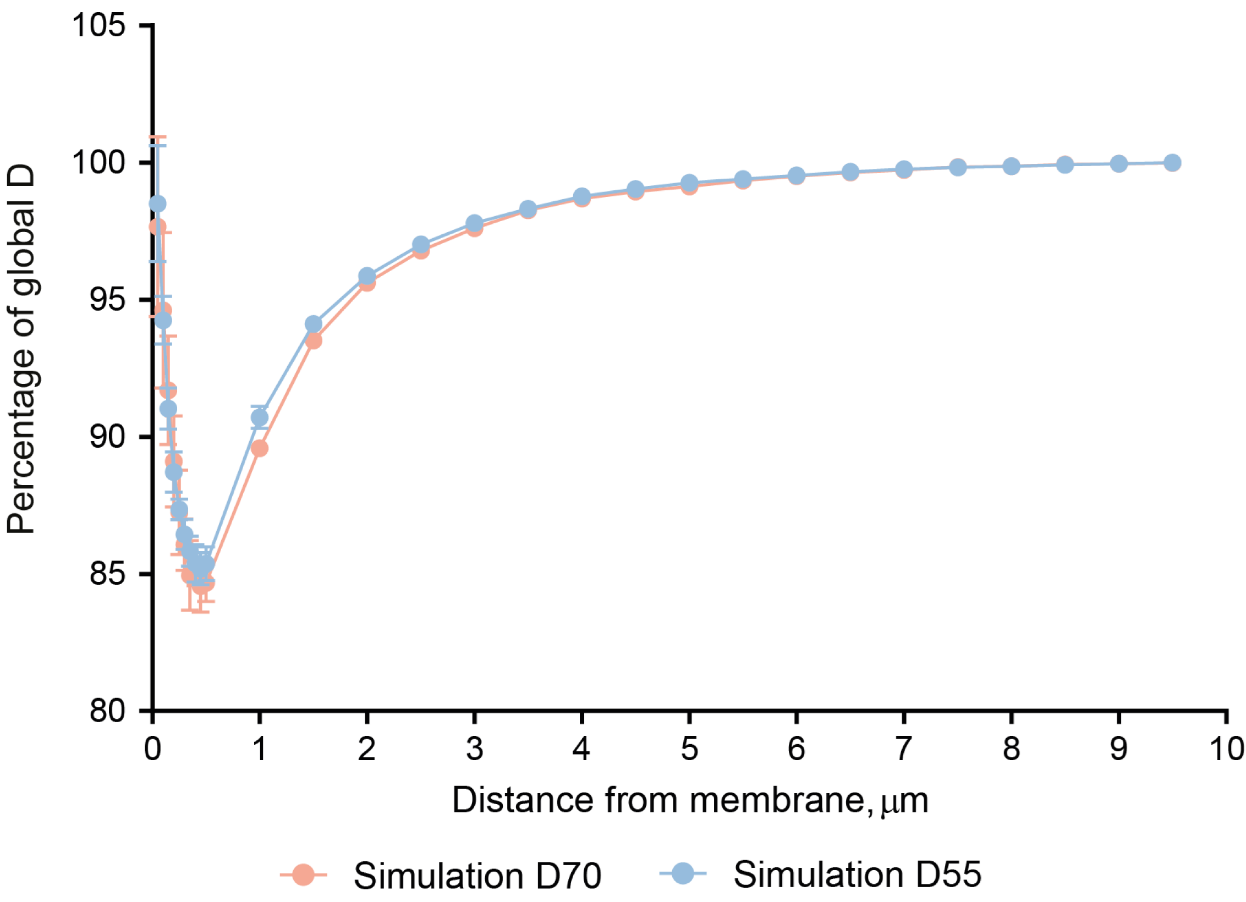

**Supplementary Figure S3. Confinement simulations using Smoldyn software^1,2^**. Diffusion of particles with input diffusion coefficients of 70 µm^2^/s (D70) and 55 µm^2^/s (D55) in a sphere with a radius of 10 µm and an elastic boundary was simulated. The minimum in the diffusion coefficient is seen at approximately 0.5 µm from the membrane. The decrease in diffusion coefficient is smaller at distances close to the membrane due to the simulated elasticity of the boundary.

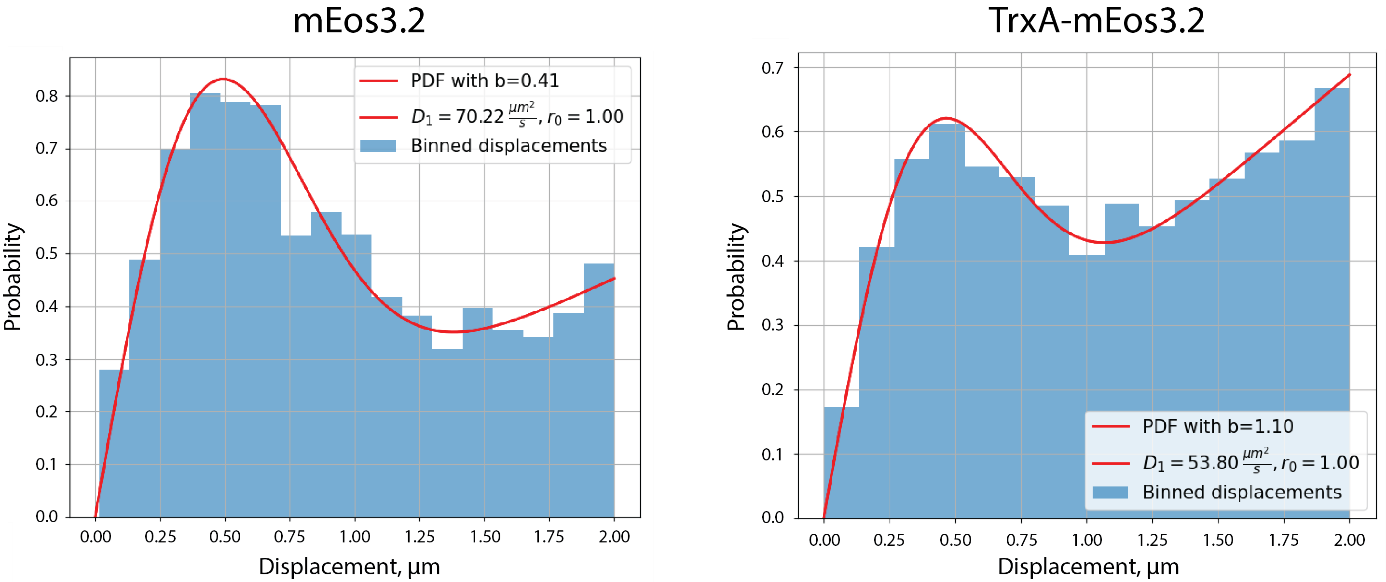

**Supplementary Figure S4. Representative probability density function fitting profiles of the mEos3.2 and TrxA-mEos3.2 displacements in DOPC GUVs.** Here, the internal medium was composed of 50 mM KPi pH 7.0, 200 mM of sucrose plus 2 µM of mEos3.2 or TrxA-mEos3.2. D_1_ and *b* refer to diffusion coefficient and background correction coefficient, respectively.

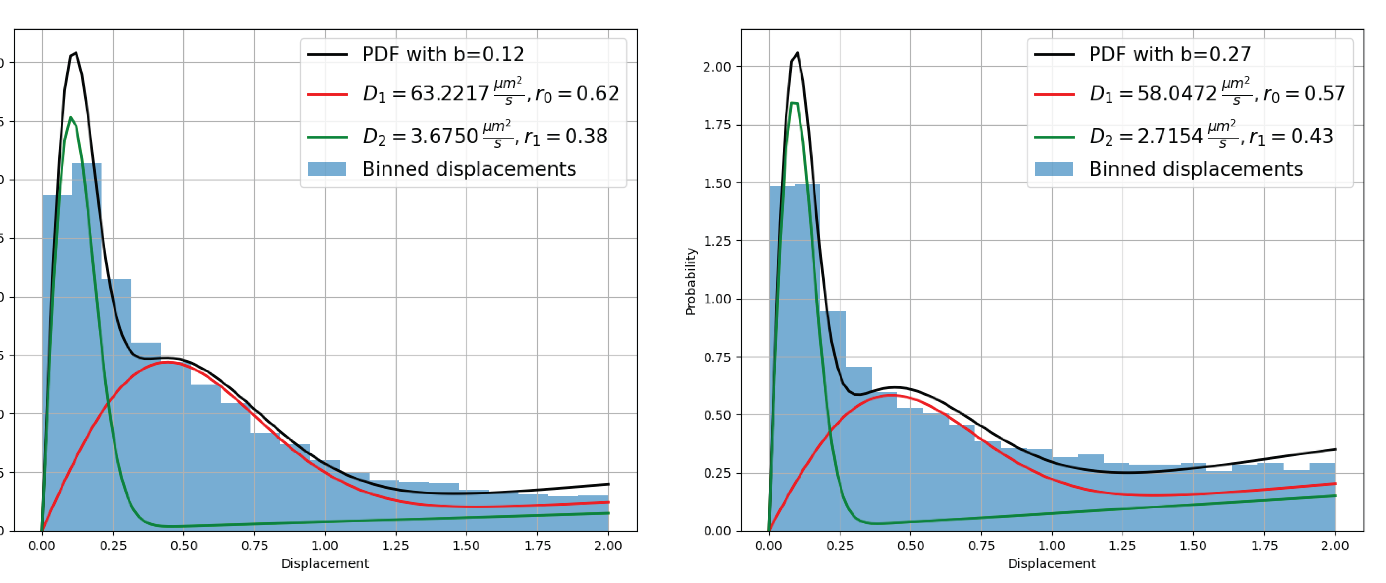

**Figure S5. Two representative probability density function fitting profiles of LactC2-mEos3.2 in DOPC:DOPS (99.99:0.001 mol%) GUVs.** Here, the internal medium was composed of 50 mM KPi pH 7.0, 200 mM of sucrose plus 0.2 µM of LactC2-mEos3.2. D_1_ and D_2_ refer to diffusion coefficient of the fast (freely diffusing) and slow (membrane bound) LactC2-mEos3.2; b corresponds to the background correction coefficient.

**Supplementary Table S1: Primers for USER cloning of LactC2-mEos3.2.** All primers contain uracil residues to enable recognition and cleavage by the USER (Uracil-Specific Excision Reagent) Enzyme, to facilitate the cloning.

| LactC2_USER_FWD | 5’-AATTAACCA**U**GAGCGCAATTAAA |
| --- | --- |
| LactC2_USER_REV | 5’-AGATCTTTTAT**U**AATGATGATGATGATGGTG |
| LactC2_Backbone_USER_rev | 5’-ATGGTTAAT**U**CCTCCTGTT |
| Lactc2_Backbone_USER_for | 5’-AATAAAAGATC**U**GCAGCTGG |

**Supplementary Table S2: Parameters of the proteins used in this work.** Parameters were determined using the ProtParam tool of Expasy. * without 10-histidine tag. Net surface charges were calculated from AlphaFold2-predicted protein structures using the **findSurfaceCharge** script in PyMOL.

| Protein | M_W_, kDa | Extinction coefficients, M^-1^ cm^-1^ | Surface charge (e) |
| --- | --- | --- | --- |
| mEos3.2 | 27.04 | 27515 | +2.21 |
| TrxA_EC_-mEos3.2 | 38.70 | 41620 | -2.48 |
| TrxA_HV_-mEos3.2 | 38.96 | 36120 | -13.47 |
| LactC2-mEos3.2 | 45.35* | 72560 | 5.02 |

**Supplementary Table S3.** **Exact diffusion coefficients measured in this work.** Data presented as mean ± SEM. Number of replicates indicated in brackets in the table. (A) Diffusion coefficients (<*D*>) corresponding to Figure 1B and GUVs diameters (d). (B) Diffusion coefficients (<*D*>) corresponding to Figure 1F and GUVs diameters (d). (C) Diffusion coefficients (<*D*>) corresponding to Figure 3A and GUVs diameters (d). (D) Diffusion coefficients corresponding to Figure 4C for fast and slow component; and percentages of corresponding diffusing species from Figure 4D. Diameter of GUVs is 15.6 ± 3.4 µm.

**A**

| Sucrose concentration | mEos3.2 | | TrxA-mEos3.2 | |
| --- | --- | --- | --- | --- |
|  | <*D*>, µm^2^/s | d, µm | <*D*>, µm^2^/s | d, µm |
| 100 mM | 80 ± 10 (13) | 21.2 ± 4.3 | 67 ± 10 (12) | 20.4 ± 3.9 |
| 200 mM | 73 ± 10 (15) | 16.9 ± 3.8 | 56 ± 8 (13) | 19.8 ± 4.0 |
| 400 mM | 63 ± 6 (8) | 19.5 ± 3.9 | 46 ± 4 (14) | 20.1 ± 4.0 |

**B**

| Lipid composition | mEos3.2 | | TrxA_EC_-mEos3.2 | | TrxA_HV_-mEos3.2 | |
| --- | --- | --- | --- | --- | --- | --- |
|  | <*D*>, µm^2^/s | d, µm | <*D*>, µm^2^/s | d, µm | <*D*>, µm^2^/s | d, µm |
| DOPC | 73 ± 10 (15) | 16.9 ± 3.8 | 56 ± 8 (13) | 19.8 ± 4.0 | 51 ± 10 (15) | 18.4 ± 3.1 |
| DOPC:DOPG | 72 ± 11 (15) | 19.8 ± 3.2 | 54 ± 4 (9) | 18.8 ± 1.5 | 53 ± 8 (14) | 18.7 ± 4.0 |
| DOPC:DOPG:DOPE | 73 ± 8 (16) | 21.7 ± 4.4 | 63 ± 6 (7) | 22.5 ± 5.5 | 61 ± 13 (9) | 17.6 ± 2.7 |

**C**

| KPi concentration | TrxA_EC_-mEos3.2 | | TrxA_HV_-mEos3.2 | |
| --- | --- | --- | --- | --- |
|  | <*D*>, µm^2^/s | d, µm | <*D*>, µm^2^/s | d, µm |
| 10 mM | 70 ± 6 (14) | 19.7 ± 2.7 | 60 ± 5 (10) | 20.6 ± 2.3 |
| 50 mM | 56 ± 8 (13) | 19.8 ± 4.0 | 51 ± 10 (15) | 18.4 ± 3.1 |
| 200 mM | 60 ± 4 (13) | 20.0 ± 3.2 | 45 ± 6 (11) | 22.2 ± 3.0 |

**D**

| Component | LactC2-mEos3.2  <D>, µm^2^/s | Percentage of |
| --- | --- | --- |
| Fast | 60 ± 7 (11) | 57 ± 12 % |
| Slow | 4.3 ± 2.0 (11) | 43 ± 12 % |

**Supplementary Table S4.** **Summary of p-values in this work.** (A) Significance values of Figure 1F. (B) Significance values of Figure 3A.

| Condition | p-value | Significance |  |
| --- | --- | --- | --- |
| mEos3.2: DOPC - DOPC:DOPG | 0.7603 | not significant | ns |
| mEos3.2: DOPC - DOPC:DOPG:DOPE | 0.8752 | not significant | ns |
| mEos3.2: DOPC:DOPG - DOPC:DOPG:DOPE | 0.6312 | not significant | ns |
| TrxA_EC_mEos3.2: DOPC - DOPC:DOPG | 0.489 | not significant | ns |
| TrxA_EC_mEos3.2: DOPC - DOPC:DOPG:DOPE | 0.0655 | not significant | ns |
| TrxA_EC_mEos3.2: DOPC:DOPG - DOPC:DOPG:DOPE | 0.0041 | very significant | ** |
| TrxA_HV_mEos3.2: DOPC - DOPC:DOPG | 0.5399 | not significant | ns |
| TrxA_HV_mEos3.2: DOPC - DOPC:DOPG:DOPE | 0.0398 | significant | * |
| TrxA_HV_mEos3.2: DOPC:DOPG - DOPC:DOPG:DOPE | 0.0772 | not significant | ns |
| Condition | p-value | Significance |  |
| TrxA_EC_mEos3.2: 10 mM KPi - 50 mM KPi | < 0.0001 | extremely significant | *** |
| TrxA_EC_mEos3.2: 10 mM KPi - 200 mM KPi | < 0.0001 | extremely significant | *** |
| TrxA_EC_mEos3.2: 50 mM KPi - 200 mM KPi | 0.136 | not significant | ns |
| TrxA_HV_mEos3.2: 10 mM KPi - 50 mM KPi | 0.0158 | significant | * |
| TrxA_HV_mEos3.2: 10 mM KPi - 200 mM KPi | < 0.0001 | extremely significant | *** |
| TrxA_HV_mEos3.2: 50 mM KPi - 200 mM KPi | 0.0931 | not significant | ns |

**Supplementary Table S5.** **Size of GUVs, used for close-to-border and “ring” areas analysis.** Number of analysed GUVs in mentioned in the brackets.

| Protein | Lipid composition | Diameter, µm |
| --- | --- | --- |
| mEos3.2 | DOPC | 14.2 ± 2.4 (4) |
|  | DOPC:DOPG | 18.6 ± 3.8 (4) |
|  | DOPC:DOPG:DOPE | 20.9 ± 4.1 (4) |
| TrxA_EC_-mEos3.2 | DOPC, 10 mM KPi | 18.4 ± 2.7 (4) |
|  | DOPC, 50 mM KPi | 17.7 ± 4.9 (4) |
|  | DOPC, 200 mM KPi | 19.5 ± 2.2 (6) |
|  | DOPC:DOPG | 19.1 ± 1.8 (5) |
|  | DOPC:DOPG:DOPE | 19.8 ± 2.0 (4) |
| TrxA_HV_-mEos3.2 | DOPC, 10 mM KPi | 19.9 ± 2.3 (7) |
|  | DOPC, 50 mM KPi | 18.9 ± 3.4 (5) |
|  | DOPC, 200 mM KPi | 21.2 ± 3.3 (6) |
|  | DOPC:DOPG | 18.6 ± 3.9 (5) |
|  | DOPC:DOPG:DOPE | 16.9 ± 2.0 (6) |
